# High rates of apoptosis visualized in the symbiont-bearing gills of deep-sea *Bathymodiolus* mussels

**DOI:** 10.1101/422840

**Authors:** Piquet Bérénice, Shillito Bruce, H. Lallier François, Duperron Sébastien, C. Andersen Ann

## Abstract

Symbiosis between *Bathymodiolus* and Gammaproteobacteria enables these deep-sea mussels to live in toxic environments like hydrothermal vents and cold seeps. The quantity of endosymbionts within the gill-bacteriocytes appears to vary according to the hosts environment. We investigated the hypothesis of a control of the endosymbionts density by apoptosis, a programmed cell death. We used fluorometric TUNEL-method and active Caspase-3-targeting antibodies to visualize and quantify apoptotic cells in mussel gills. To avoid artefacts due to depressurization upon specimen recovery from the deep-sea, we compared the apoptotic rates between mussels recovered unpressurised, versus mussels recovered in a pressure-maintaining device, in two species from hydrothermal vents on the Mid-Atlantic Ridge: *Bathymodiolus azoricus* and *B. puteoserpentis.* Our results show that pressurized recovery had no significant effect on the apoptotic rate in the gill filaments. Apoptotic levels were highest in the ciliated zone and in the circulating hemocytes, compared to the bacteriocyte zone. Apoptotic gill-cells in *B.* aff. *boomerang* from the pockmarks off the Gulf of Guinea, show similar distribution patterns. Deep-sea symbiotic mussels have much higher rates of apoptosis in their gills than the coastal mussel *Mytilus edulis* without chemolithoautotrophic symbionts. We discuss how apoptosis might be one of the mechanisms that contribute to the adaptation of deep-sea mussels to toxic environments and/or to symbiosis.

## Introduction

Symbiosis is of major significance to life on Earth. Because symbiosis is a mixture of cooperation and conflict between two (or several) partners, each of them must initiate and carry on a continued dialogue, and control their partners’ interaction. Many symbioses involve partners from different domains of life, such as a eukaryote host and a bacterial symbiont. This questions how a reciprocal control can occur between so distantly related organisms, and whether some of the mechanisms involved might be universal. One of the mechanisms by which an animal host can control populations of its symbionts is apoptosis. Apoptosis is a programmed cell death involving three main steps: (1) nuclear condensation and fragmentation, (2) cell-wall budding into apoptotic bodies, and (3) their release and possible phagocytosis by neighboring cells [1, 2]. Apoptosis plays multiple roles in normal cell turnover, during development, and in the immune system [3]. Its role in symbiosis is also documented: in the cereal weevil *Sitophilus* for example, apoptosis participates to regulate the densities of the endosymbiotic bacterium *Sodalis pierantonius.* The symbiont provides essential amino acids that allow the host to rapidly build its protective exoskeleton. Endosymbionts multiply in young adults, but when the cuticle is built, the symbionts are rapidly eliminated through apoptosis of the host cells that contain them [4]. In corals, bleaching occurs in response to heat stress, when the host releases dinoflagellate symbionts through apoptosis and autophagy linked in a see-saw manner, such that when apoptosis is inhibited autophagy is initiated as a back-up mechanism, and vice-versa [5]. In addition apoptosis might act as a post-phagocytic winnowing mechanism in the symbiotic system [6].

At deep-sea hydrothermal vents and cold seeps, the animals that dominate in terms of biomass live in association with chemosynthetic bacteria, which sustain most of their nutrition [7-10], but the role apoptosis could play in regulating symbiosis has barely been explored. In the vestimentiferan tubeworms *Riftia pachyptila*, living at vents, and *Lamellibrachia luymesi* from cold seeps, the sulfur-oxidizing symbionts are located within cells of the trophosome. These cells differentiate and proliferate from the trophosome lobule center, then migrate towards the periphery of the lobule where they undergo apoptosis [11]. Ultrastructural observations at the periphery of the trophosome lobules show that the symbionts are digested in vacuoles leading to extensive myelin bodies, then remnants of symbionts disappear, while apoptotic nuclei with clumped chromatin patches occur [11]. Thus, in *Riftia* as in the weevil, apoptosis appears to be involved in the process of symbiont regulation to recover the metabolic investment from the symbiotic phase.

Deep-sea mussels house very dense populations of endosymbionts inside specialized gill epithelial cells, the bacteriocytes, [12, 13]. In fact, *Bathymodiolus* constitute by far the densest microbial habitats, although they usually host a very limited diversity of symbiont lineages [7, 8]. The relevance to the topic of symbiont control lies in the fact that the association is particularly flexible, with abundances of their symbionts (Sulfur- and/or methane-oxidizers) that can vary within hours depending on the availability of symbiont substrates in the surrounding water [8, 14-18]. The symbionts also rapidly disappear if their substrates are absent [17-19]. Ultrastructural studies of the gill cells have indicated intracellular digestion of the symbionts within lysosomes as an important carbon transfer mechanism [20-22] suggesting that the host can access the energy stored in its symbionts by killing and digesting them (i.e. a process compared to “farming”). Enzymatic studies involving the detection of acid phosphatases have shown that some energy from the symbionts can also be transferred to the host through molecules leaking from live symbionts (i.e. a process compared to “milking”) [23]. Recent results from whole-gill tissue transcriptomic analyses in the vent species *Bathymodiolus thermophilus* indicated that high symbiont loads are correlated with underexpression of the genes inhibiting apoptosis, suggesting that when the symbionts are abundant, apoptosis might be activated [24, 25]. It can thus be hypothesized that apoptosis is a mechanism by which the hosts control the number of symbionts inside their gills, and possibly obtain their carbon. High throughput sequencing and transcriptomic analyses have shown the great importance of the apoptotic signaling pathways in the gill tissue of several species of *Bathymodiolus* [26-28].

Apoptosis in mollusks is generally triggered by the interaction between immune cells and parasites or pathogens [29]. The high degree of evolutionary conservation of biochemical signaling and executing pathways of apoptosis indicate that programmed cell death likely plays a crucial role in homeostasis and functioning of molluscan immune system [29-31]. Apoptosis is particularly complex in mollusks, and caspases are key molecules that are activated in two major apoptotic pathways: the extrinsic or death receptor pathway, and the intrinsic or mitochondrial pathway [31]. The executioner caspase-3 activates a heterodimer protein, the DNA fragmentation factor that is responsible for the completion of DNA fragmentation [31]. However alternative caspase-independent pathways have also been evidenced in mollusks, and cross-talk between different pathways might also be involved [30, 31]. Apoptosis was shown to be induced in *Bathymodiolus azoricus* in response to *Vibrio diabolicus* exposure [32]. However the immune gene responses in *B. azoricus* appeared tied to the presence of endosymbiont bacteria, as the progressive weakening of its host transcriptional activity correlates with the gradual disappearance of endosymbiont bacteria from the gill tissues during the extended acclimatization in the sulfide and methane-free aquaria [32]. Thus, the presence of symbionts might modulate the apoptotic patterns in deep-sea symbiotic mussels compared to their coastal mussel relatives without chemolithoautotrophic symbiont.

While several studies have highlighted an important activity of apoptotic signaling factors in three species of *Bathymodiolus (B. azoricus, B. manusensis* from hydrothermal vents, and *B. platifrons* from cold seeps (Bettencourt et al., 2010; Wong et al., 2015; Zheng et al., 2017)), transcriptomic studies do not give a visual account of this apoptotic activity within the tissues. The present study is the first microscopic investigation of the general distribution patterns of apoptotic cells in the gills, and the first attempt to quantify apoptosis in the different gill cell types of *Bathymodiolus.* We chose to focus on three species, namely *Bathymodiolus azoricus, B. puteoserpentis* and *B.* aff. *boomerang.* The first two often dominate the macrofauna at Mid-Atlantic Ridge (MAR) hydrothermal vent sites [33, 34]. The third occurs at cold seeps located on the continental margin in the Gulf of Guinea [35]. The former two species are phylogenetically sister species and are also closely related to *B. boomerang* [12]. All three species harbor methane- and sulfur-oxidizing bacteria that co-exist within host cells. Their gills comprise several cell types: epidermal ciliated cells and mucous goblet cells, bacteriocytes hosting the symbionts, interspaced by intercalary cells, and finally circulating hemocytes [36]. Our aim was to investigate, whether apoptosis might contribute to regulating symbiont densities. We thus tested, whether apoptosis preferentially occurs in bacteriocytes, in particular the most densely populated ones.

As the recovery of deep-sea specimens from several thousand meters depth usually involves a depressurization stress that might alter gene expression and disturb the cell machinery, leading to artefacts [37, 38] we performed a comparative analysis of apoptosis in specimens recovered with and without depressurization upon collection. Specimens of *B. azoricus* and *B. puteoserpentis* were recovered using an hyperbaric chamber (PERISCOP - Projet d’Enceinte de Remontée Isobare Servant la Capture d’Organismes Profonds) that allows maintaining their native pressure and temperature throughout recovery, avoiding recovery bias [39]. Control specimens recovered without PERISCOP were included. The cold seep *B.* aff. *boomerang* was analyzed to identify potential seep versus vent habitat-related differences, and *Mytilus edulis* was used as a non-chemolithoautotrophic-symbiont-bearing control mussel. Apoptosis was visualized in gill tissue sections by the TUNEL method (Transferase dUTP Nick-End Labeling) [11, 40, 41], a common and standard way of visualizing the fragmented DNA in the nucleus of cells undergoing apoptosis. To further support that observed patterns actually correspond to apoptosis, we performed immunolocalization of active Caspase-3, a form of this enzyme that is the overall convergent node of molecular cascades leading to irreversible apoptosis in mollusks [31]. Altogether, this study provides to our knowledge the first visual overview of apoptosis in relation to symbiosis in chemosynthetic mussels.

## Materials and Methods

### Specimen collection

This analysis was conducted on 40 individuals of *Bathymodiolus* (Bivalvia, Mytilidae) consisting in 21 *Bathymodiolus azoricus* and ten *B. puteoserpentis* from hydrothermal vents, and nine *B.* aff. *boomerang* from cold seeps. Specimens of *B. azoricus* were collected during the BioBaz 2013 cruise on the Mid-Atlantic Ridge [42]: ten specimens were sampled from the vent fields of Menez Gwen (MG2 marker: 37°50.669N; 31°31.156W, 830m depth) and eleven specimens from Rainbow site (France 5 marker: 37°17.349N; 32°16.536W, at 2270 m depth). *B. puteoserpentis* were sampled during the BICOSE 2014 cruise [43]. All ten specimens were sampled on the vent site Snake Pit, close to the “Elan” marker (23°22’54”N, 44°55’48”W, at 3520 m depth). For these two species, half of the specimens were recovered in a standard (i.e. unpressurized) way, and the other half using the PERISCOP hyperbaric vessel. The standard collection was done in a waterproof sealed box (BIOBOX) containing local deep-sea water, in which the mussels were exposed to a depressurization corresponding to the depth of their habitat during recovery (approx. 0.1 MPa every 10 m). The pressurized recovery was performed using a small device named the “Croco” sampling cell [44], which was transferred into the PERISCOP pressure-maintaining device [39]. PERISCOP was released from the seafloor through a shuttle system, and surfaced within 45 to 100 minutes. Pressure was monitored during ascent with an autonomous pressure sensor (SP2T4000, NKE Instruments, France). To compare apoptosis between vent and seep mussels, nine specimens of *Bathymodiolus* aff. *boomerang* were collected in a standard manner from the Régab pockmark site in the Gulf of Guinea (M1 marker 5°47.89S, 9°42.62E, 3150 m depth and M2 marker 5°47.85S, 9°42.67E, 3150 m depth) during the 2011 cruise WACS [45]. All three cruises were aboard the RV *Pourquoi Pas?* using the ROV *Victor 6000.* In addition to deep-sea mussels, nine coastal mussels *(Mytilus edulis)* in two sets were analyzed. First set were wild *M. edulis* collected on rocks from the intertidal seashore in front of Roscoff Marine Station in January 2014 (n=4 individuals analyzed). The second set of *M. edulis* were cultivated mussels bought from the fishmongers in Paris in January 2017 (n= 5 individuals analyzed). All data concerning the specimens are shown in the Supplementary Table 1.

### Sample fixation, inclusion and FISH experiments

Mussel gills were dissected at 4°C, within 10 minutes after recovery (releasing pressure of the PERISCOP in the case of isobaric recoveries). Maximum shell length, height under the umbo, and width of the closed shell were measured with a caliper (see supplementary figure 1). Anterior and posterior gill fragments were fixed in 4% formaldehyde in sterile-filtered seawater (SFS) for 2 hours. Gills were then rinsed in SFS, and dehydrated in increasing series of ethanol (50, 70 and 80%, 15 min each). In the laboratory, gills were embedded in polyethylene glycol (PEG) distearate: 1-hexadecanol (9:1), cut into 8 μm-thick sections using a microtome (Thermo, Germany), recovered on SuperFrost Plus slides (VWR International, USA), and stored at −20°C. Fluorescence *in situ* hybridization (FISH) experiments were performed to confirm the localization of symbionts following the protocol and using probes described previously [15]. The gill lamellae were sectioned ventrally, so that each filament appeared in cross-section with its frontal end facing externally towards mantle and shell, and its abfrontal end facing the inner gill lamellae towards the foot. (See supplementary figure 2).

**Figure 1:**
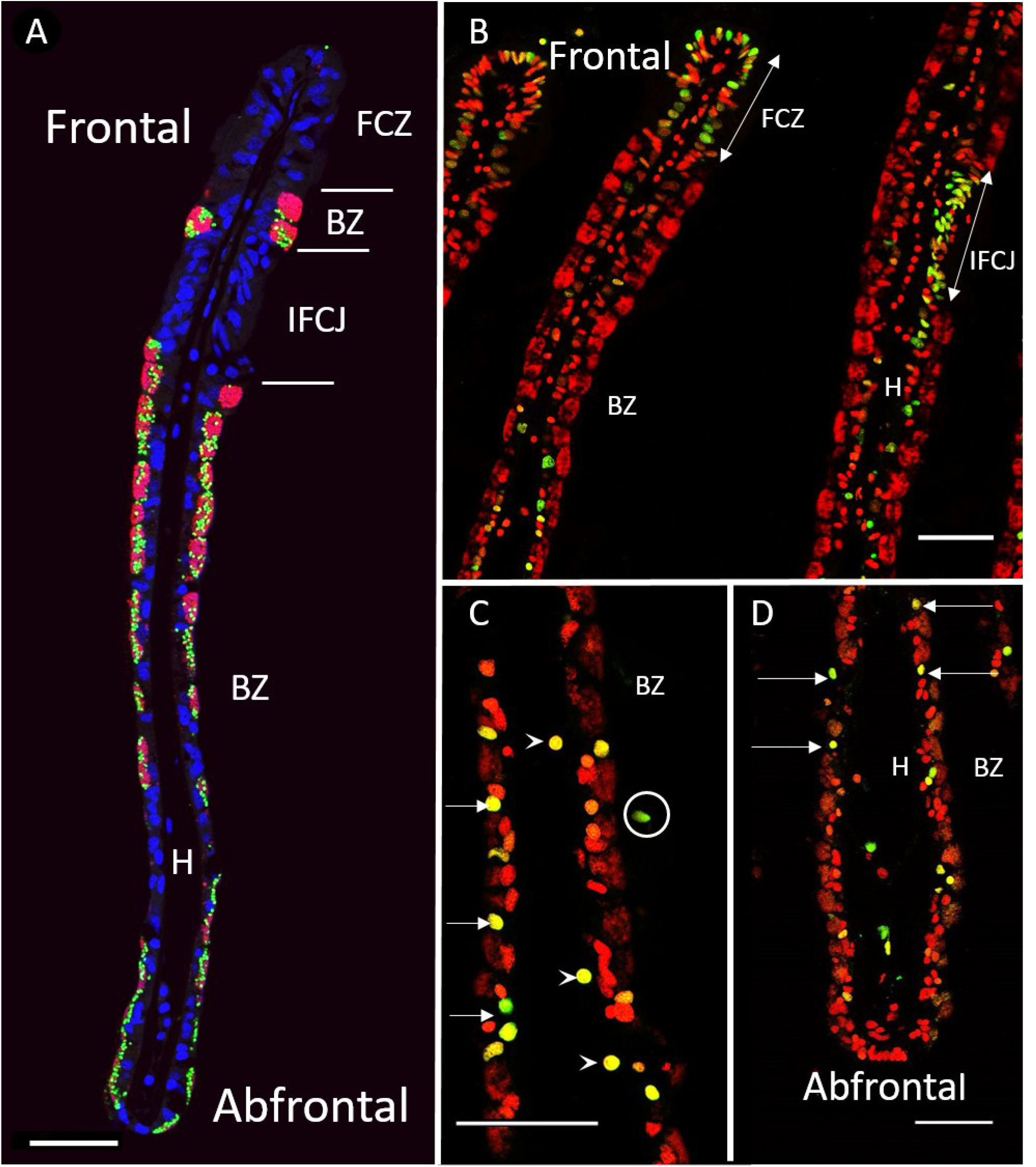
FISH and TUNEL labelling on gill filaments of *B. azoricus.* A: Transverse section of gill filaments with FISH labelling. Nuclei from host tissue in blue (DAPI); sulfur-oxidizing symbionts in pink, and methane-oxidizers in green. B, C and D TUNEL-labelled nuclei in green, DAPI staining in red. Arrows point to apoptotic nuclei. B: Ciliated zones often contain many labelled nuclei. C: Bacteriocyte zone that displays many apoptotic nuclei (arrows), apoptotic hemocytes (arrowheads) and an apoptotic cell detached from the basal lamina (circle). D: Abfrontal zone of the gill filament showing apoptotic nuclei in cells with no symbiont (arrows), and apoptotic cells detached from the basal lamina (circles). Abbreviations: BZ: Bacteriocyte zone; FCZ: Frontal Ciliated Zone; H: Hemolymph zone; IFCJ Inter-Filamentary Ciliated Junction. Scale bars: 50μm.

**Figure 2:**
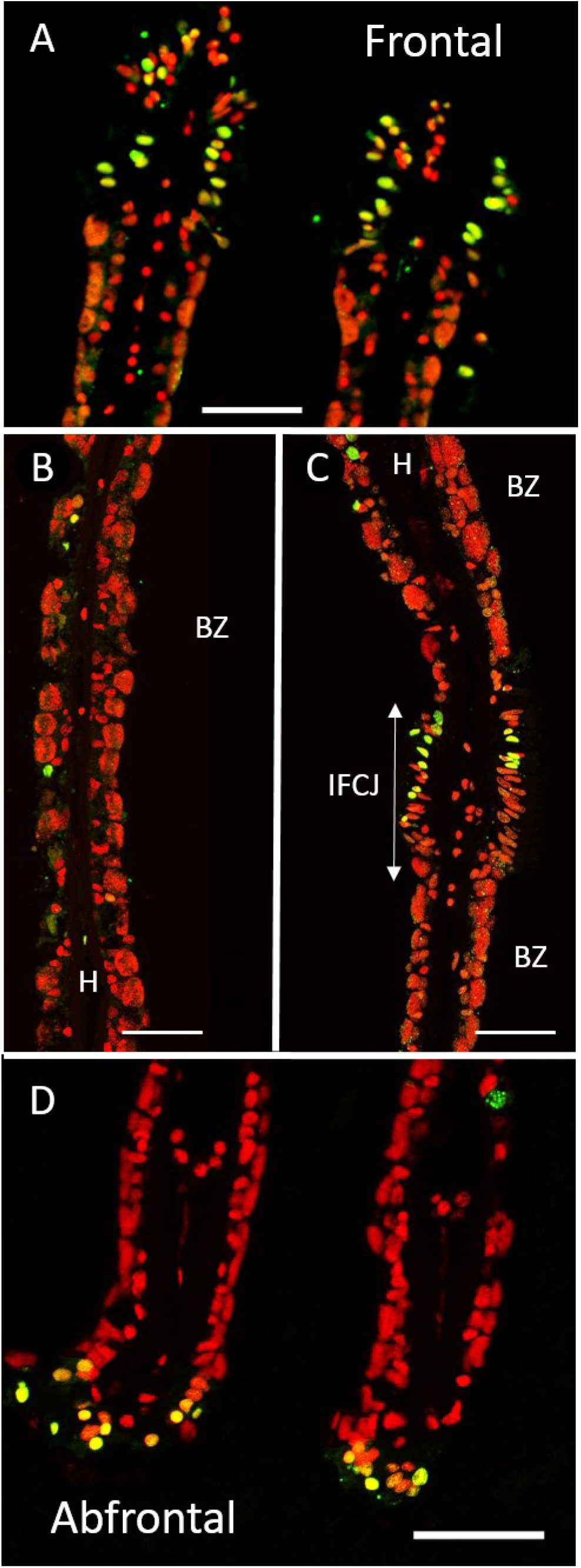
TUNEL labelling on gill filaments of *B. puteoserpentis.* TUNEL-labelled nuclei in green, DAPI staining in red. A: Frontal ciliated zones often contain many labelled nuclei. B: Bacteriocyte zone that displays only a few apoptotic nuclei. C: Inter Filamentary ciliated Junction often contain many labelled nuclei. D: Abfrontal zone of the gill filament with apoptotic nuclei. Abbreviations: BZ: Bacteriocyte zone; FCZ: Frontal Ciliated Zone; H: Hemolymph zone; IFCJ Inter-Filamentary Ciliated Junction. Scale bars: 50μm.

### TUNEL (Transferase dUTP Nick-End Labeling)

For the detection and quantification of apoptosis, we used the TUNEL method with the *in-situ* cell death detection kit following manufacturer’s instructions (ROCHE, Germany). All slides were first dewaxed then rehydrated by immersion in decreasing series of ethanol. Tissues were rinsed with PBS 1 X and permeabilized by proteinase K (20μg.ml^-1^) to enable the binding of dUTP. Various incubation times were tested, and 8 min gave the best result. Slides were incubated at 37°C for 1h30 with fluorophore dUTP and enzyme rTdT. Slides were rinsed in 3 PBS bathes (10 min each) to remove the unfixed fluorescein-12-dUTP. In a last step, DNA of all nuclei was labelled with 4’6’-Diamidino-2-phenylindole (DAPI) that is incorporated in the SlowFade^®^ Gold Antifade Mounting reagent (Invitrogen, USA). For each analyzed individual, positive and negative controls were performed on adjacent serial sections. The positive control involved a pre-incubation with DNase I (3U/ml, 10 min) before running the full protocol. It provokes artificial fragmentation of DNA and exposes 3’OH ends, which bind the green fluorochrome and leads to all nuclei being labelled. The negative control follows the same protocol, but omitting the rTdT enzyme. It enables to reveal autofluorescence of the tissues and unspecific fixation of the fluorophore.

### Immunolocalization of active Caspase-3

Slides were dewaxed and rehydrated as for the TUNEL experiments. They were then covered in blocking solution (PBS 10X, 2% BSA, 0.3% triton) for 2 h at 4°C. Tissue sections were incubated with the rabbit polyclonal active anti-Caspase-3 primary antibody (directed against a peptide from the P17 fragment of the activated human/mouse Caspase-3) (R&D system, USA). This primary incubation lasted 2 hours at 4°C. Slides were rinsed in PBS 10X three times and covered with the secondary antibody: Alexa Fluor 488-labelled goat anti-rabbit (Invitrogen, dilution 1:500) for 1 h at room temperature. After 3 baths in PBS 10X, slides were mounted with SlowFade^®^. Negative controls were obtained by omitting the primary antibody.

### Image acquisition and analysis

Slides were observed under a BX61 epifluorescence microscope (Olympus, Japan), and pictures were taken under an SP5 confocal (Leica, Germany) at the x40 magnification. This magnification enabled a clear resolution to count each individual labeled nucleus. Entire cross-sections of the TUNEL-labelled gill filaments were covered in full in about 3 to 4 pictures (figures 1 and 2). The exposure time (or gain on the confocal) was standardized and identical for all pictures. The 3D-acquisitions were obtained by acquiring images every 0.5 μm throughout the thickness of the section. Slides were observed at 505 nm and 555 nm wavelengths (TUNEL and Caspase 3 signals, respectively), and 400 nm (DAPI). Classical immuno-localization are shown with all labeling in their original color (DAPI in blue), but for TUNEL labelled nuclei we had to change the DAPI color into red. Indeed blue DNA-labelling overlaid by green (i.e. apoptosis TUNEL labeling) leads to a complex mixture of various levels of green and blue, often making quantitative interpretations difficult. To enable an easy quantitative analysis of the TUNEL images, we chose to visualize DAPI in red, so that a double labelling would lead to a third distinct color (yellowish). Thus, a TUNEL-labelled nucleus appears bright green when its amount of fragmented DNA is great (i.e. when a nucleus is in an advanced state of apoptosis), but yellow or orange, when its amount of fragmented DNA is respectively low or very low (i.e. when a nucleus is in an early state of apoptosis). At first, we counted green and yellowish nuclei separately, but as they both expressed apoptosis, and early and advanced states were randomly distributed in the gill lamellae (figures 1 and 2), we finally pooled them together.

Images were analyzed by counting the labelled nuclei using the free Image J software [46]. However care was taken to count separately each respective cell-type: hemocytes, ciliated cells and finally bacteriocytes and intercalary cells (the two latter being close neighboring cell types, hard to distinguish under the fluorescence microscope, they were counted together) (figure 1 and 2). Thus for each image, percentages of TUNEL-labelled nuclei were obtained from the hemolymph zone (HZ), the ciliated zone (CZ), and the bacteriocyte zone (BZ) by calculating the ratio between the number of TUNEL-labelled nuclei and the number of DAPI-labeled nuclei present in each respective zone. A fourth value, the total percentage for the whole picture, was obtained by summing all labeled nuclei (whatever their cell-type) on the total number of DAPI-stained nuclei in a given picture.

### Comparisons between species, zones and treatments

The percentage of TUNEL nuclei labelled was used for all analyses after an Arcsine transformation [47]. The normality of datasets was tested (Shapiro-Wilk test), which revealed a non-normal distribution of the data. Non-parametric tests were thus applied for inter-groups comparisons (Mann-Whitney-Wilcoxon and Kruskal-Wallis) with the Bonferroni correction for multiple comparisons. All statistical analyses and graph plots were performed using R (R Development Core Team, version 3.3.3).

## Results

All analyses were performed on transverse sections of the gill lamellae (supplementary figure 2). From the FISH results, it is noteworthy that bacteriocytes close to the frontal ciliated zone contained large quantities of endosymbionts, and that both the height of the bacteriocytes and their symbiont density decreased towards the abfrontal zone in *B. puteoserpentis*, and *B. azoricus* (figure 1A). This frontal/abfrontal decrease in symbiont density also appears with the DAPI staining, as this labels not only the host nuclei, but also the DNA of the bacterial symbionts.

## Visualization of apoptosis in Bathymodiolus

The distribution of TUNEL positive cells in *B. azoricus* and *B. puteoserpentis* are shown in Fig. 1 and 2, respectively. In negative controls (supplementary figure 3B), no fluorescence at all was observed compared to positive controls (supplementary figure 3A), except a bright auto-fluorescence signal seen in clustered round granules. These granules are thought to correspond to mucus droplets present in goblet-cells interspersed among the ciliated cells and along the epithelium of the gill filaments. This autofluorescence signal was easy to distinguish from TUNEL signal, as the latter was typically much smaller and brighter. Nuclei labelled in the TUNEL experiments were more or less bright. As TUNEL labels the free 3’OH fragments, we assume that the strongly labelled nuclei showed a higher amount of DNA fragments compared to the weakly stained ones. Since apoptosis is a progressive process occurring within a few hours, we hypothesize that the former were in a more advanced apoptotic state. Both weak (orange or even yellow) and strong-labelled nuclei (green) were thus counted together to estimate the percentage of apoptotic nuclei in the various cell-types of the gill lamellae (ciliated zone, bacteriocyte zone and hemolymph) (figure 1and 2).

**Figure 3:**
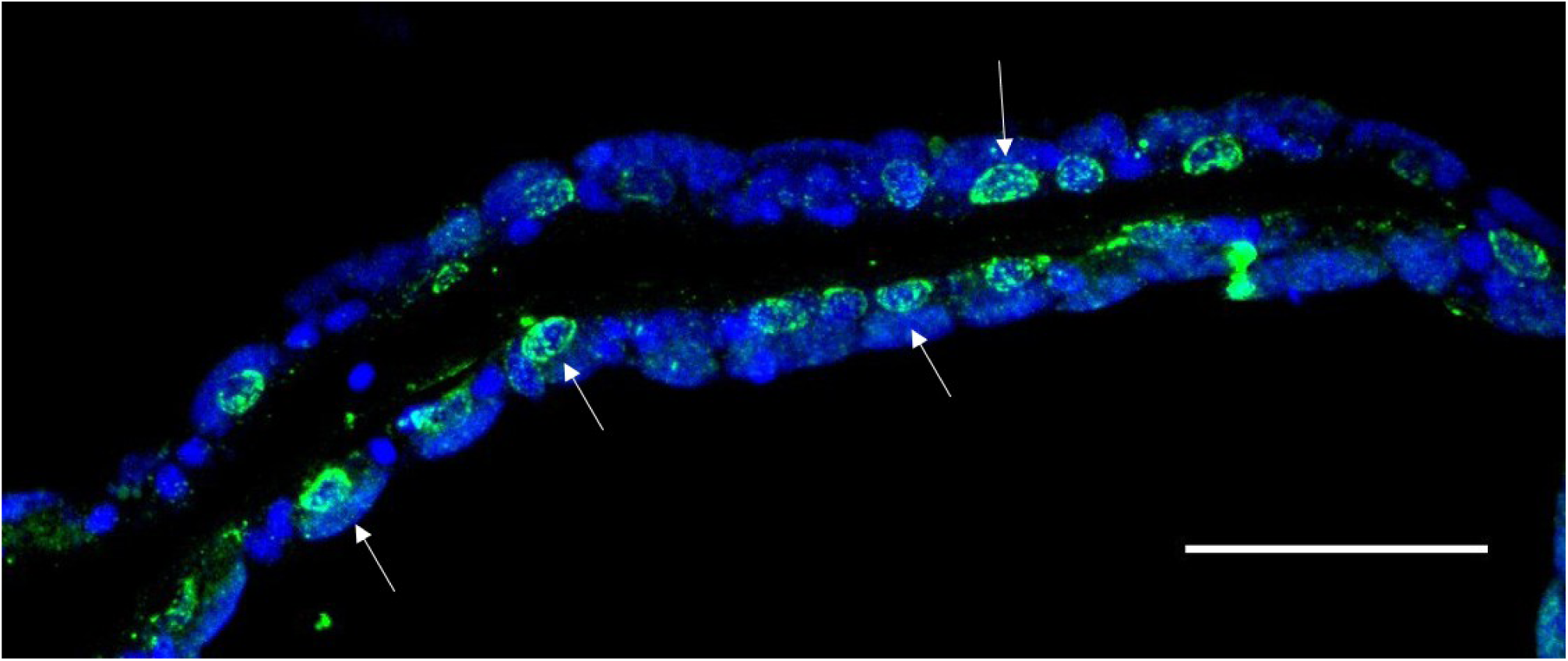
Active Caspase-3 labelling in the gill of *B. azoricus.* Typical labelling in green surrounds the nucleus (arrows). DAPI isignal in blue. Scale bar: 50μm.

The ciliated frontal zone often is the brightest labelled zone with many strongly labelled cells: up to one cell out of two in some filaments (Fig. 1B, 2A) giving a clustered appearance of the apoptotic cells in the frontal ciliated zone. However the closest neighbor filament may sometimes only have a few or no labelled cells at all, indicating a great spatial variability in the tissue taken all together. Hemocytes were present on all images, but in varying numbers. Many hemocytes are visible on figure 1C for example, compared to figure 1B, but more of them were labelled in the latter. TUNEL-labelling in the bacteriocyte zone was heterogeneous: very few nuclei were labelled close to the frontal ciliated zone (figure 1B and figure 2A), and then increasingly more were labelled when reaching the abfrontal zone (figure 1D and figure 2D). In the abfrontal zone, the bacteriocytes were most of the time devoid of bacteria. Sometimes, a bacteriocyte seemed to be lacking in the abfrontal zone, leaving a “hole” in the epithelial tissue (figure 1C-D and figure 2 B-D).

Active Caspase 3 immunohistochemistry assays yielded strong signals within the cytoplasm of cells, and particularly concentrated around the nuclei (figure 3). On serial sections within the same individual, the distribution patterns of active Caspase-3 were similar to the TUNEL signal observed, confirming that TUNEL actually revealed apoptotic cells. In general, the active Caspase-3 antibody seemed to label more nuclei than TUNEL.

### Patterns of apoptosis in Bathymodiolus azoricus and B. puteoserpentis

In total, 31 individuals from the two species were analyzed, representing a total of 19,612 DAPI-labelled nuclei counted on 206 images. The specimens came from deep-sea sites located at different depths, namely Menez Gwen (−830 m), Rainbow (−2270 m) and Snake Pit (−3520 m). Sampled specimens were either recovered unpressurised (non-isobaric) or with a pressurized (isobaric) recovery. In the latter, PERISCOP reached the surface while retaining 83.6, 76.5 and 70.4% of the native deep-sea pressure at the Menez Gwen, Rainbow and Snake Pit sites, respectively.

First of all, comparing the global counts in gills of all individuals from all three sites, we noted that there is a great variation in the percentage of apoptotic nuclei among individuals (see supplementary figure 4 A-C). This strong heterogeneity indicates that results of statistical analyses must be treated with caution. In the ciliated (CZ) and bacteriocyte zones (BZ), the percentage of TUNEL-labelled nuclei was not significantly different among the three MAR sites and the two recovery types (CZ: Kruskal-Wallis test chisquared =6.07, df=5, p-value=0.30; BZ: Kruskal-Wallis test chi-squared=3.69, df=5, p-value=0.60) (See figure 4 for BZ and supplementary figure 5 A for CZ). Similarly, the hemolymph zone showed no significant difference among sites and recovery types (HZ: Kruskal-Wallis test chi-squared=10.74, df=5, p-value=0.05678) (supplementary figure 5 B). These quantitative results confirm the visual patterns and intensities of TUNEL-labelling in gill filaments of mussels that did not reveal any self-evident difference between specimens from the three sites in either recovery type. For all specimens put together, quantifications indicated median values of 41.3% (±27%) and 34.5% (±31%) labelled nuclei in the ciliated and hemolymph zones, respectively. In comparison, the bacteriocyte zone displayed only 19.3% (±24%) of nuclei that were labelled, (figure 5) which is significantly less than in the other two (Kruskal-Wallis test chisquared=27.639, df=2, p-value<0.001, post-hoc test with Bonferroni correction between HZ and BZ, p-value=0.0002; between CZ and BZ, p-value <0.0001).

**Figure 4:**
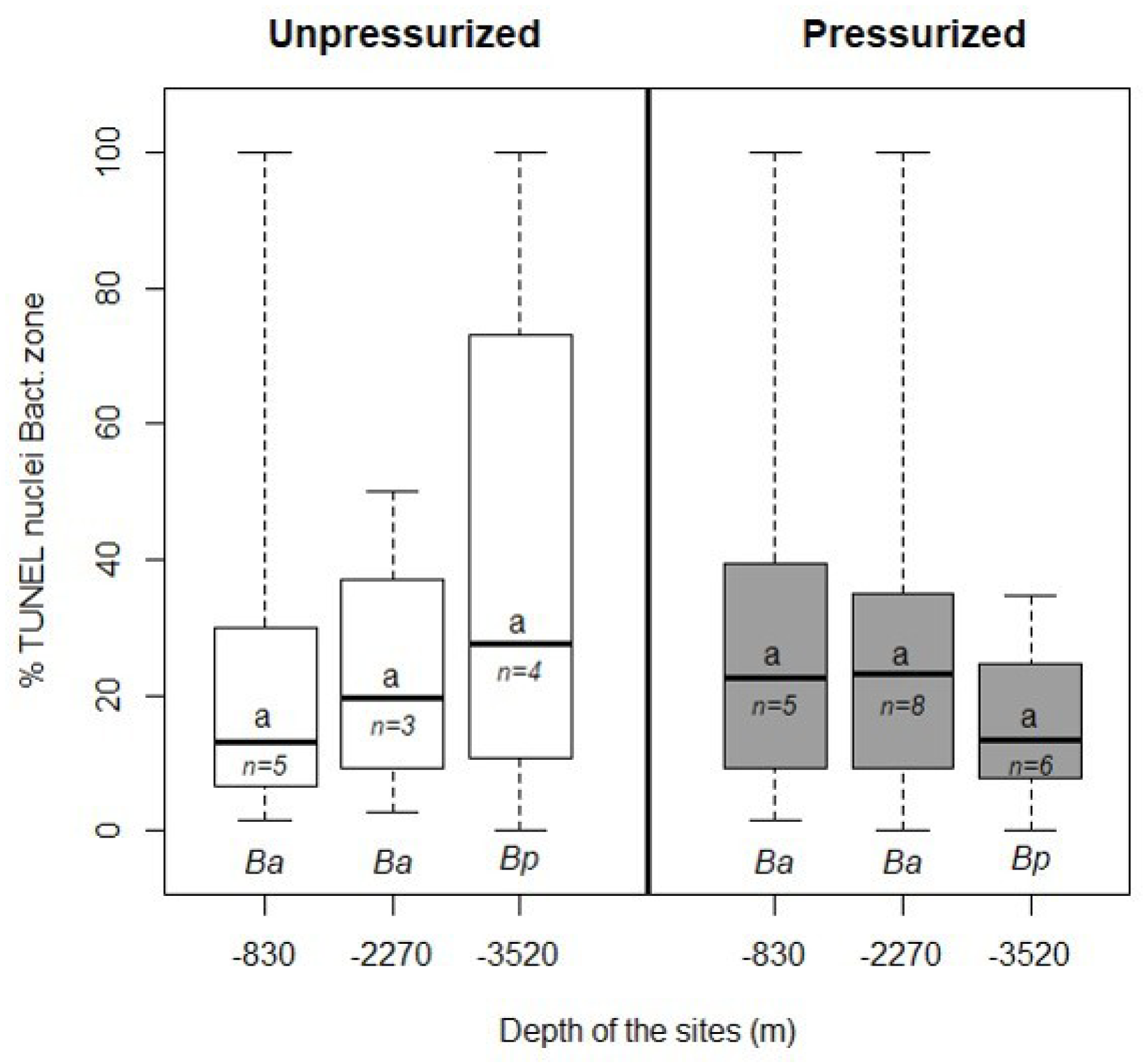
Percentages of apoptotic nuclei in the bacteriocyte zone of *B. azoricus* (Ba) and *B. puteoserpentis* (Bp) from the three sites. White boxplots indicate specimens from non-isobaric, and grey from isobaric recoveries, respectively. Percentages were ‘not significantly different (a) in pairwise Wilcoxon test with Bonferroni’s standard correction. The letter *n* indicates the number of individuals analyzed for each species. Boxplot whiskers correspond to minimal and maximal values on a single image, the line inside the box is the median, and the upper and lower frames of the boxes represent the first and third quartile respectively.

**Figure 5:**
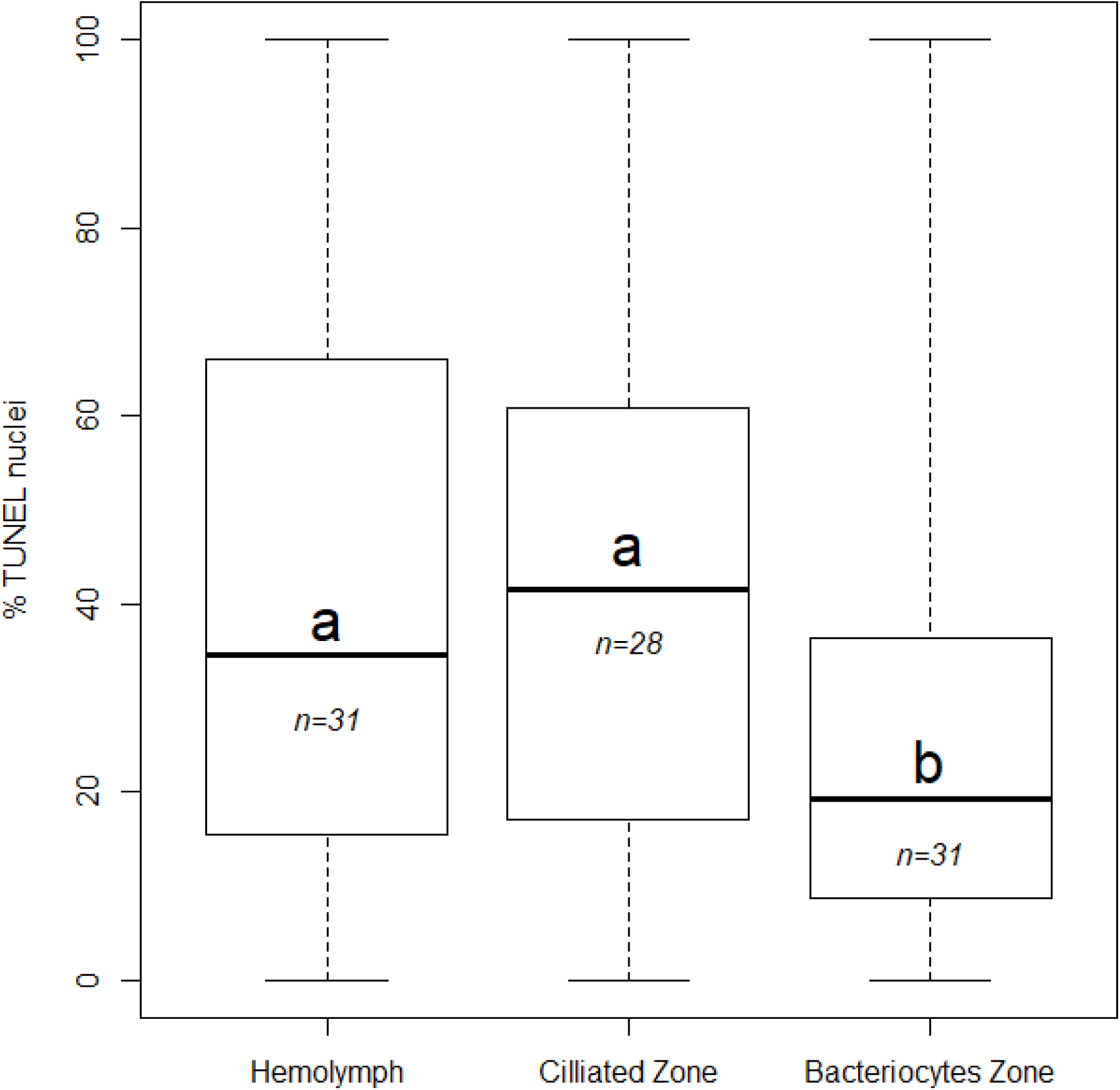
Percentages of apoptotic nuclei in the hemolymph, ciliated and bacteriocyte zones of *B. azoricus* and *B. puteoserpentis.* Different letters (a, b) indicate the plots that are statistically different (Wilcoxon test) and n indicates the number of specimens. Boxplot whiskers indicate minimal and maximal values on a single image, the line inside the box is the median, and the upper and lower frames of the boxes represent the first and third quartile respectively.

### Patterns of apoptosis in the seep species Bathymodiolus aff. boomerang

TUNEL labelling was applied to the gill filaments of *Bathymodiolus* aff. *boomerang* 9 specimens, 40 pictures representing 3,424 nuclei) (figure 6 A). Percentages were significantly different between zones (Kruskal-wallis test: chi-squared=25.00, df=2, p-value<0.0001). As in the two studied vent species, apoptosis was more frequent in the ciliated zone (median: 36%, ±23%) and then in the hemocytes zone (median: 21% ±9%), compared to the bacteriocyte zone (median: 9% ±23%). Overall these results suggest similar patterns in Mytilidae from hydrothermal vents and cold seeps (figure 7). Visually, the density of bacteria inside bacteriocytes seems to be more constant between bacteriocytes located in the frontal and in the abfrontal distal edge in *B.* aff. *boomerang* compared to the two vent mussels.

**Figure 6:**
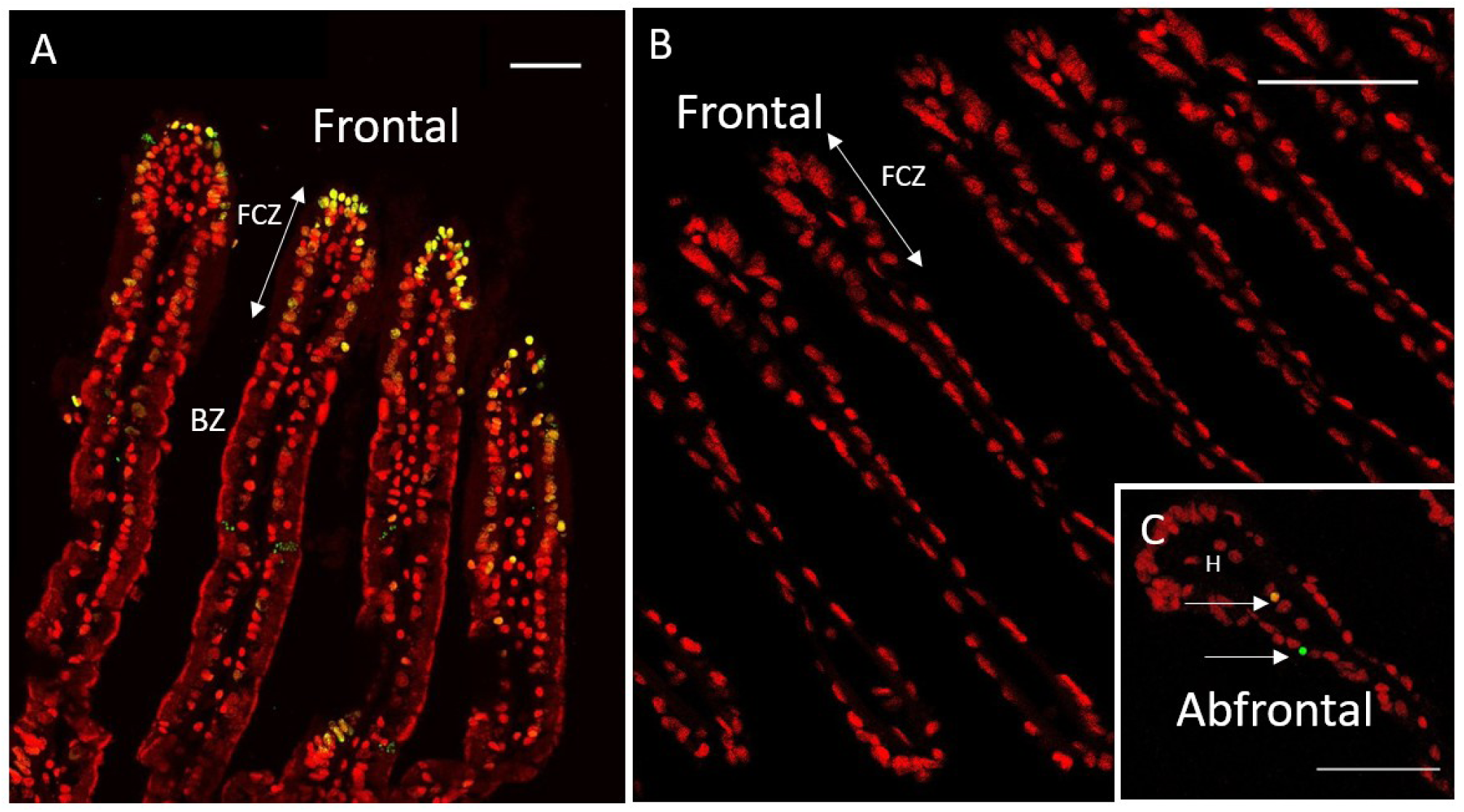
TUNEL labelling on gill filaments of *Bathymodiolus* aff. *boomerang* (A) and *Mytilus edulis* (B-C). TUNEL-labelling in green or yellow, DAPI staining in red. A: *Bathymodiolus* aff. *boomerang* displays many labelled nuclei mostly in the frontal ciliated zone. B-C: *Mytilus edulis* gills show very few TUNEL-labelled nuclei, two are visible in the insert (C). Same abbreviations; as in figure 1. Scale bar: 50μm.

**Figure 7:**
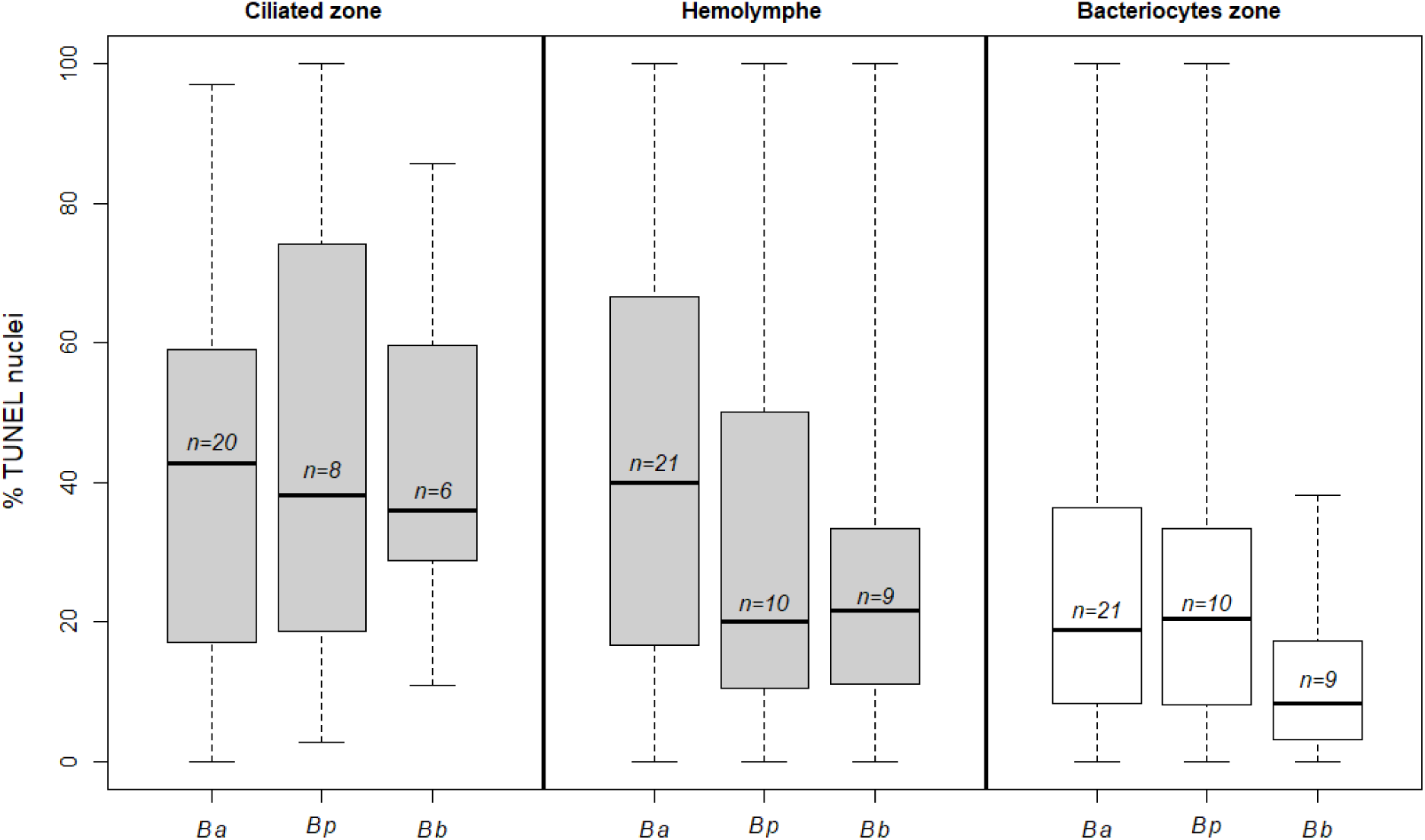
Percentages of apoptotic nuclei compared between vent and seep species in the ciliated, hemolymph and bacteriocyte zones of *B. azoricus, B. puteoserpentis* and *B.* aff *boomerang* (Ba, Bp and Bb, respectively). The letter *n* indicates the number of specimens. Boxplot whiskers indicate minimal and maximal values on a single image, line inside the box is the median, the line inside the box is the median, and the upper and lower frames of the boxes represent the first and third quartile respectively.

### Patterns of apoptosis in the gill of coastal Mytilus edulis

Gill tissues of ***Mytilus edulis*** were labelled by the TUNEL method (9 specimens from two different localizations, 28 images, 4,091 nuclei). Both the wild mussels, collected on the intertidal rocks near the marine station in January and the commercial mussels bought in Paris in January, gave the same distribution pattern of apoptotic cells. The number of labelled nuclei was very low in comparison to vent and seep mussels (figure 6B) (median: 1.6%, ±1.6%) and nearly the same percentage between pictures. In *M. edulis*, the majority of labelled nuclei were hemocytes (median: 6.9%, ±10.9%) compared to the epidermal cells (median: 0.9%, ±1.4%).

## Discussion

### Relevance of TUNEL labelling to the study of apoptosis in deep-sea mussels

During apoptosis DNA undergoes fragmentation, and the TUNEL assay labels the free 3’OH ends that are generated. However, other mechanisms can lead to DNA fragmentation that would result in a positive TUNEL labelling, and for this reason the TUNEL methodology has been criticized [48-50]. It is nevertheless a standard procedure that has already been applied in mollusk tissues, and in association with active caspase-3 immunolocalization to detect apoptosis in the chemosymbiotic bivalve *Codakia orbiculata* [40, 51]. Caspase-3 exists within the cytosol as inactive dimers. All apoptotic pathways in mollusks converge towards a step of cleavage of this zymogen that ultimately results in the triggering of apoptosis [31, 52]. Although it may have other roles, the activation of Caspase-3 usually leads to cell death by apoptosis. Because other caspases activate necrosis and necroptosis pathways, the occurrence of active Caspase-3 signals rather supports that most observed TUNEL signals actually correspond to apoptotic nuclei [53]. It should however be kept in mind that another apoptotic pathway exists that is Caspase-independent, as it can take place without activating Caspase-3 [54, 55].

### Deep-sea mussels display high rates of apoptosis in their gills

Recovery stress has long been cited as a factor that may prevent accurate assessment of the physiology of deep-sea organisms [39, 56, 57]. In this study, we found no statistical difference between percentages of apoptotic cells between gills from specimens of *Bathymodiolus azoricus* and *B. puteoserpentis* recovered classically (non-isobaric) and those recovered using the isobaric sampling cell PERISCOP. Thus depressurization during recovery does not seem to trigger massive apoptosis in the gills, despite that depressurization, by disrupting cells, may alter the distribution of different actors of apoptosis that pass through different cell compartments (mitochondria, nucleus, cell membrane) [39]. No differences were observed between hydrothermal vent sites, whatever their depth, further suggesting that apoptosis is a natural phenomenon in the gills of deep-sea mussels. Estimates of rates of apoptosis based on TUNEL labelling were much higher in the gills of deep-sea mussels including the seep species *B.* aff. *boomerang*, compared to their coastal relative *Mytilus edulis* lacking chemolithotrophic symbionts. This comparison has been made with two different sets of *Mytilus:* wild and cultivated, yielding the same scarce TUNEL-distribution pattern. At this stage it is not possible to ascertain, whether the much higher levels measured in *Bathymodiolus* are linked with their deep-sea chemosynthetic habitats (for example due to the abundance of xenobiotic compounds and oxidative stressors), with the occurrence of symbionts, or with their different evolutionary history.

### Patterns and potential roles of apoptosis in deep-sea mussels

A major finding in this study is that rates of apoptosis vary considerably among *Bathymodiolus* individual specimens, contrary to *Mytilus*, for a similar investigation effort. This could reveal distinct physiological status among the *Bathymodiolus* at the time of the analysis. Nonetheless, it should be underlined that apoptosis is known to be a rapidly occurring mechanism, in a time range of only a few hours [1]. The distribution of apoptotic cells in a tissue often shows a clustered pattern, as for instance in themammalian intestine, where apoptotic signal appears typically confined to a cluster of neighboring villi, while other areas of the mucosa appear unstained [41]. A sporadic distribution might ensure a local recycling of the organ, without harming its global function, and might indeed be a characteristic of the apoptotic phenomenon. In the case of *Bathymodiolus*, gills often showed a few TUNEL-positive cells in some crosssections, and even clusters in the frontal ciliated area of some filaments, but none in the frontal zone of neighboring filaments. This clustered pattern may also contribute to the observed heterogeneity among our individuals. Anyway, the great inter-individual variation prevents from drawing any definitive conclusion, even based on statistical tests of which the results have to be treated with caution. Nevertheless, a clear trend is that the rates measured in the bacteriocyte zone are lower than in the ciliated and hemocyte zones. The very high rates of apoptosis in the ciliated gill cells, in particular compared to *M. edulis*, are intriguing because these cells do not contain chemolithotrophic symbionts. The gills of *Bathymodiolus* are clearly different from those of the coastal mussels for which, at any given size, the gill surface area is around 20 times smaller [8]. *Bathymodiolus* gills are particularly thick and over-developed, which might lead to a higher metabolic demand in order to properly circulate water and vital compounds. Ciliated cells also display very abundant mitochondria [8, 13, 58]. One hypothesis could thus be that the great metabolic activity and numerous mitochondria produce large amounts of Reactive Oxygen Species (ROS), resulting in oxidative stress that may lead to increased apoptosis [59]. A possible adaptation would be to increase the turnover rates of the ciliated cells. To achieve this, high rates of cell proliferation could be expected, and this has to be tested. Another hypothesis is linked with fluid toxicity and the direction of water flow in the gills. Ciliated cells are indeed the first exposed to the surrounding fluid, and thus exposed to the highest levels of reduced compounds including toxic sulfur and various metals that possibly trigger apoptosis [60]. This may also explain why symbionts tend to be more abundant in the bacteriocytes that are close to the regions that are more exposed to their substrates (mainly methane and sulfide), since these compounds might be more easily accessible to symbionts first in the frontal zone compared to the abfrontal zone, sustaining bacterial growth. Among deleterious compounds, copper ions were for example recently hypothesized to cause cell apoptosis in *Bathymodiolus azoricus.* Cadmium was shown to cause apoptosis on isolated *Crassostrea gigas* hemocytes after 24h *in vitro* incubation in the range of 10-100 μmol.L^-1^ [61]. Gills of mussels are known to accumulate metals, and *Bathymodiolus azoricus* exposed to cadmium display antioxidant enzymatic activities that may be partly due to the endosymbionts. These could contribute to protect bacteriocytes, but not the symbiont-free ciliated cells, resulting in higher apoptosis rates in the latter [62]. *In vivo* incubation experiments at *in situ* pressure would be necessary to test these oxidative stress and fluid toxicity hypotheses.

Apoptosis of circulating and resident hemocytes have a high baseline level in mollusks [29], and we estimated that 34.5 % of the hemocytes were apoptotic in *B. azoricus* and *B. puteoserpentis*. The immune system of mussels relies on innate immunity only, and hemocytes play a key role in the immune response. They are also suspected to have a role in detoxification [58]. In *Crassostrea virginica*, the most common apoptotic cell type is the hemocyte. The apoptotic index for *Crassostrea* hemocytes, calculated the same way as herein, but using another visualization method (Apoptag), ranges between 23 and 99% with a mean of 46% [63]. In another study, *Crassostrea virginica* displayed between 4.5% and 15.3% of apoptotic circulating hemocytes (Annexin-V assay) [64]. These results are congruent with the rates observed in this study, and could reveal overall high apoptosis rates in the hemocytes of bivalves, visually confirming the importance of apoptosis in the innate immune system.

Despite the fact that apoptosis rates are high in *Bathymodiolus* bacteriocytes compared to those in *M. edulis* epithelial gill-cells, they are only roughly half of those measured in ciliated cells and hemocytes. Results from transcriptomic studies and their subsequent interpretation by other authors led to the hypothesis that apoptosis was the main mechanism that allowed mussels (initially *Bathymodiolus thermophilus*) to control the amount of symbionts in their gills. The underlying idea was that bacteria-filled bacteriocytes would undergo apoptosis in a way similar to that observed in the tubeworm *Riftia pachyptila*, in which this mechanism participates to the recovery of symbionts carbon, and recycling of the animal cells [11]. On the contrary, our observations show that the bacteriocytes that contain only a few, or no bacteria at all, are mainly those that enter into apoptosis. Lower levels of apoptosis in bacteriocytes, in particular in those where symbionts are the most abundant, could be explained by the bacterial protection concerning the toxic compounds, which could provoke apoptosis of the cell. This could explain why cells with fewer bacteria, mostly located in the distal (i.e. abfrontal) edge away from the ciliated zone and possibly most depleted in symbiont substrates, would preferentially enter apoptosis, although this is merely hypothetical at this stage. In bacteriocytes near the ciliated zone, bacteria are most probably digested [21, 23, 65].

Overall, it appears that high rates of apoptosis are a normal feature in deep-sea mussel gills physiology. Rather than being a direct mechanism used to kill the most bacteria-rich cells in order to gain carbon, apoptosis seems to be involved in the overall dynamics of the gill organ itself. Indeed, observed patterns indicate that the cells harboring few to no bacteria are more often apoptotic. This can be interpreted in the light of gill specialization to symbiosis, with over-developed gills requiring stronger cilia activity, which possibly leads to higher turnover of ciliated cells, and habitat characteristics including the presence of toxic compounds. Apoptosis mechanisms are known to play major roles in other symbioses [4, 5, 11, 66]. For example *Codakia orbiculata*, a shallow water bivalve that hosts Gammaproteobacteria in its gills can lose and reacquire symbionts. After loss of the symbionts, the bacteriocytes multiply massively to reacquire bacteria, while the cells in excess are eliminated by apoptosis [40]. A similar mechanism may be occurring in mussels. At first, apoptosis could appear as a heavy cost for the host, especially given that it may endanger the whole organ. However, neighboring cells could phagocytose the cells that undergo apoptosis, allowing recycling of their constituents. Apoptotic cells also tend to detach from the basal lamina, and once detached they may be treated as food particles by the mussels, again allowing recycling. The correlation observed in previous transcriptome analyses [25] between apoptotic rates and overall symbiont content may then be indirect: hosts with more bacteria tend to have larger gills [8], and thus more active ciliated cells, so overall more cells in apoptosis. High rates of apoptosis should thus be regarded as an integral part of the adaptation of deep-sea mussels to their symbioses and their habitats, and visualization pattern indicates that apoptosis probably plays more complex roles than previously assumed. This study has only investigated adult specimens, but because mussels acquire their symbionts in their post-larval stage and throughout the host lifetime [67, 68], the coupling of symbiont acquisition with apoptosis should be explored at all developmental stages. Further study should also investigate the impact of stress (toxic compounds, temperature) and those of symbiont loss (when *Bathymodiolus* is moved away from the fluids compounds [18, 69]) on the rates of apoptosis, as well as cell proliferation patterns. Because of the relative simplicity of the mussel holobiont, its high bacterial load and high levels of apoptosis, deep-sea mussels may prove useful models to further investigate the links between apoptosis, autophagy and symbiosis.

## Acknowledgements

We thank the captain and crew on board the Resarch Vessel *“Pourquoi pas?”* and the ROV *“Victor 6000*” (Ifremer) for their invaluable contributions to this work. A special thanks to chief scientists Dr. Karine Olu (WACS 2011), Pr. François Lallier (BioBaz 2013) and Dr. Marie-Anne Cambon-Bonavita and Dr. Magali Zbinden (BICOSE 2014). We are indebted to Dr. Kamil Szafranski for his help with the deep-sea mussels *B. azoricus* and *B. puteoserpentis* recovered on board during the BICOSE and BioBaz cruises. We are grateful to Antoine Mangin, Myriam Lebon and Marie Louvigné for their technical assistance as undergraduate students in the lab. This research was supported by Institut Universitaire de France project ACSYMB, Université Pierre et Marie Curie and ARED Région Bretagne project FlexSyBi (contract number 9127, grant to BP), BALIST program (ANR-08-BLAN-0252), EU projects EXOCET/D (FP6-GOCE-CT-2003-505342), MIDAS (FP7/2007-2013, grant agreement n° 603418) and CNRS and IFREMER for cruise funding. We are grateful to the microscopy platform Merimage (Roscoff, France) partner core facilities and to the Institute of Biology Paris-Seine Imaging Facility, supported by the “Conseil Regional Ile-de France”, CNRS and Sorbonne Université.

## Author’s contribution

B.P did the histology work, the microscopy imaging, the statistical analyses and wrote a first version of the paper. B.S. performed the isobaric mussel collections with the PERISCOP pressure-maintaining device during the two cruises (BioBaz 2013 and BICOSE 2014). F.L. enabled the collections of deep-sea mussel during the BioBaz 2013 cruise. S.D fixed the mussels during the WACS 2011 and BioBaz 2013 cruises and supervised BP’s analyses in the AMEX-team in Paris. AA fixed the *M. edulis* and contributed to BP’s histology work in the ABICE-team in Roscoff. S.D and AA are the collaborative designers of the study. S.D., A.A. and B.P. wrote the latest version of the paper, all authors have read and corrected its final draft.

